# The orexigenic force of olfactory palatable food cues in sated rats

**DOI:** 10.1101/2021.07.06.451251

**Authors:** Fiona Peris-Sampedro, Iris Stoltenborg, Marie V. Le May, Pol Solé-Navais, Roger A. H. Adan, Suzanne L. Dickson

**Affiliations:** Department of Physiology/Endocrine, Institute of Neuroscience and Physiology, The Sahlgrenska Academy at the University of Gothenburg, Gothenburg, Sweden; Department of Obstetrics and Gynaecology, The Sahlgrenska Academy at the University of Gothenburg, Gothenburg, Sweden; Brain Center Rudolf Magnus, Department of Translational Neuroscience, University Medical Center Utrecht, Utrecht University, The Netherlands

**Keywords:** Food cues, Feeding, Ghrelin, Arcuate nucleus, AgRP, POMC

## Abstract

**Background:** Environmental cues recalling palatable foods are ubiquitous and motivate eating beyond metabolic need, yet the timing of this response and whether it can develop towards a non-palatable readily available food remain elusive. Although there is increasing evidence indicating that external stimuli in the olfactory modality can communicate with the major hub in the feeding neurocircuitry, the hypothalamic arcuate nucleus (Arc), the identity of hypothalamic substrates has been only partially uncovered.

**Methods:** Using a palatable home-cage hidden-food paradigm, we investigate the ability of olfactory food cues to promote chow overconsumption in sated male rats, together with their impact on meal pattern. We likewise explore, by means of an immediate early gene marker, the neural mechanisms involved, including the possible engagement of the orexigenic ghrelin system.

**Results:** Olfactory detection of a familiar palatable food shifts diurnal patterns towards an increase in meal frequency to cause persistent overconsumption of chow in sated conditions. In line with the orexigenic response observed, sensing the palatable food in the environment stimulates food-seeking and risk-taking behavior, and also triggers release of active ghrelin. Olfactory food cues recruit intermingled populations of cells embedded within the feeding circuitry within the Arc, including, notably, those containing the ghrelin receptor, even when food is not available for consumption.

**Conclusions:** These data demonstrate leverage of ubiquitous food cues, not only for palatable food-searching, but also to powerfully drive food consumption in ways that resonate with heightened hunger, for which the orexigenic ghrelin system is implicated.

## Introduction

Food-linked cues in our environment are powerful triggers that encourage us to search for and consume food, even when sated (1, 2). Most studies exploring the mechanisms underpinning cue-potentiated feeding use classical Pavlovian conditioning models, in which animals learn through association that a non-food stimulus (e.g., either discrete [a light or a tone] or contextual cues) signals food availability (3–9). However, in nature, animals/humans mostly rely on conditioned cues provided by or directly derived from the food itself (e.g., smell or sight) to enhance food consumption.

The sense of smell is a central driver of food-seeking, appetite, and food preference in vertebrates, including humans (10, 11). Classical neuroanatomical and electrophysiological studies indicate the existence of an olfactory–hypothalamic axis of potential relevance for feeding control (12), and connections between the main olfactory bulb and the hypothalamic arcuate nucleus (Arc; a core hub in the feeding neurocircuitry) have been described (13, 14). Recent advances using real-time fiber photometry recordings demonstrate the engagement of orexigenic agouti-related protein (AgRP) and anorectic proopiomelanocortin (POMC) neurons, key feeding regulators residing in the Arc, in almost instantly sensing food availability in the olfactory modality prior to its consumption (15). However, an important unanswered question is whether there are other relevant cell populations in this area that respond to olfactory food cues.

As is the case for other brain areas involved in feeding control, olfactory circuits express receptors for various circulating metabolic hormones. One prominent example is the receptor for the orexigenic hormone ghrelin (growth hormone secretagogue receptor, GHSR) (11, 16). Indeed, the ghrelin system emerges as a possible link between olfactory cue-driven appetite and heightened food intake. Moreover, the possibility exists that external orexigenic signals originating from food cues in the environment (e.g., smell of food) and from intrinsic hunger signals (e.g., ghrelin) converge on overlapping circuits to impact on feeding behaviors – not only food-seeking and intake, but also, as previously demonstrated, on pathways involved in exploratory sniffing and olfactory sensitivity (17).

In this study, we sought to discover whether a chronic exposure to an olfactory palatable food-evoking cue can increase the intake of a less palatable but readily available food, such as regular chow, in sated rats subjected to a palatable home-cage hidden-food paradigm (18). Animals were exposed to an olfactory cue-rich environment recalling the palatable food whilst having access exclusively to chow. Because the timing of the cue-potentiated feeding effect, if any, remains elusive, we likewise sought to determine whether it is transient, or whether it persists, and for how long, upon cue presentation. To gain an understanding of the degree of crosstalk between internal metabolic and external olfactory food-evoking signals in sated conditions, we explored the molecular identity of neurons in the Arc that are activated upon exposure to this hidden-food paradigm. We hypothesized that ghrelin-responsive neurons are activated upon detection of the palatable food in the environment, even when food, whatever its nature, is not available for consumption.

## Methods and materials

Further detailed information about the animals, experimental procedures, and statistics used are given in Supplemental Methods.

### Animals

Single-housed adult male Sprague-Dawley rats and C57BL/6N mice were used. We used a behavioral preparation firstly developed in our lab (18), in which a metallic, perforated, opaque tea-strainer ball was suspended from each animal’s cage lid prior to single housing. During experiments, this device contained the inaccessible palatable food (peanut butter, PB; Skippy, Hormel foods, Austin, MN, USA). All experiments were approved by the local ethics committee for animal care in Gothenburg (ethics #132–2016) and complied with European guidelines (Decree 86/609/EEC).

### Feeding response and meal patterns in an olfactory PB cue-enriched environment

An automated feeding monitoring system (TSE LabMaster, Project 4261, TSE Systems, Bad Homburg, Germany) was used to analyze diurnal feeding patterns (measuring cumulative chow intake, meal frequency, meal size and ingestion rate) in an olfactory PB cue-enriched environment in both PB taste-naïve (n=16) and PB taste-familiar (n=16) rats. After PB familiarization and baseline measurements, open PB-filled tubes were encased in the perforated balls immediately after the light cycle onset and left there for 24 hr, during which odor cue-induced chow consumption was recorded. Cumulative intake of chow within 24 hr following cue removal was also recorded. To determine whether olfactory detection of PB likewise influences chow intake in mice, a cohort of PB taste-familiar mice was used in a parallel study.

### Food-seeking in familiar and risky environments enriched with an olfactory PB cue

We explored ways to quantify the dramatic food-seeking behavior observed especially in PB taste-familiar rats upon introduction of the olfactory PB cue in the home cage, which persists for around 30 min (Movie S1). PB taste-naïve (n=15) and PB taste-familiar (n=20) rats were used. Following acoustic measurements (19) of the first 15-min interaction with the perforated ball in the familiar home cage, rats from each cohort were subdivided into four subgroups (PB taste-naïve/PB cue (n=7), PB taste-naïve/No cue (n=8), PB taste-familiar/PB cue (n=10) and PB taste-familiar/No cue (n=10)) and subjected to a PB-baited open field test (adapted from (20)), which was used as a conflict-based approach to assess risk-taking behavior in food-seeking.

### Cell activation in the arcuate nucleus upon olfactory detection of PB

PB taste-naïve (n=15) and PB taste-familiar (n=13) rats were used to explore whether the olfactory PB cue activated cells in the Arc in sated conditions. *Ad libitum-*fed rats were exposed to either an olfactory PB cue-enriched environment (PB taste-naïve/PB cue, n=8; PB taste-familiar/PB cue, n=8) or to a non-enriched environment (PB taste-naïve/No cue, n=7; PB taste-familiar/No cue, n=5) for 20 min. Food access was withheld from cue exposure until sacrifice. At 80 min from cue removal, rats were anesthetized, perfused and brains harvested as described in Supplemental Methods. Free-floating brain sections were processed for immunohistochemical detection of Fos, which was used as a marker of neuronal activation (21).

### Neurochemical identification of the cells activated by the olfactory PB cue

Candidate Arc cells activated during exposure to the olfactory PB cue included those whose known distribution (based on the Allen Brain Atlas) shows some overlap with the Fos+ cells and that are known to be embedded withing the feeding circuits (AgRP, POMC and dopamine) and/or that contain GHSR (15, 22, 23). Triple fluorescent *in situ* hybridization using RNAscope® was performed to study the co-expression of Fos with GHSR-, AgRP-, POMC- and dopamine (*Tyrosine hydroxylase [Th]* probed)-containing cells in the Arc of PB taste-familiar rats (n=3-4) upon a 20-min exposure to PB odor. To this end, three independent assays were run: (i) *c-Fos*, *Ghsr* and *Agrp*; (ii) *c-Fos*, *Ghsr* and *Pomc*; and (iii) *c-Fos*, *Ghsr* and *Th*. For details refer to (21) and Supplemental Methods.

### Assessment of active ghrelin levels upon olfactory detection of PB

Another cohort of PB taste-familiar rats (n=21) was used to confirm the cue-induced hyperphagic effect. The same rats were re-exposed to either an olfactory PB cue-enriched environment (n=13) or to a non-enriched environment (perforated balls with empty tubes; n=8) for 1 hr, after which they were anesthetized and sacrificed to obtain blood for active ghrelin measurement, as described in Supplemental Methods. Food was only withheld during the 1-hr exposure to the olfactory cue.

### Statistics

The program IBM SPSS Statistics 27 (IBM Corp., Armonk, NY, USA) was used for statistical analyses. Comparisons between and within groups, and across time, were carried out by analysis of variance (ANOVA) or paired *t*-tests (see details in Supplemental Methods). Given that we did not contemplate an explicit comparison between PB taste-naïve and familiar rats, additional paired samples *t*-tests or one-way ANOVAs were used to follow up significant main effects and/or interactions.

Scatterplots and boxplots were generated using R (version 3.6.3, ggplot2 package) and other plots express the mean ± standard error. Statistical significance was set at *p* < 0.05, and values 0.05 ≤ *p* < 0.1 were considered evidence of statistical trends. Detailed statistical result annotations can be found in Supplemental Results.

## Results

### Olfactory detection of PB increases meal frequency to cause chow overconsumption in sated PB taste-familiar rats

To determine whether olfactory detection of PB can trigger overconsumption of chow in sated rats, PB taste-naïve and PB taste-familiar rats were subjected to automatic feeding recordings in the presence of PB odor (Figure 1A). Strikingly, PB taste-familiar rats showed a sustained overconsumption of chow (at a similar within-meal ingestion rate to that of the baseline recording) in the presence of the olfactory PB cue that lasted up to 16 hr post-cue introduction in the home cage (Figure 1B). Cumulative 24-hr chow intake following cue removal did not differ from that of the baseline and the cue settings (Figure S1). Notably, rats that had never tasted PB before did not change their food intake despite the presence of PB odor, suggesting that PB did not serve as a food cue in this group (Figure 1B). Meal pattern analyses revealed that PB taste-naïve rats consumed fewer meals throughout exposure to PB odor, which might reflect a hyponeophagic effect linked to the introduction of a novel odor in the cage, but compensated their lower meal frequency by increasing meal size overall (Figure 1C–D). PB taste-familiar rats, however, ate more frequently when exposed to the olfactory cue at all time points studied (Figure 1C). Interestingly, such an increase in meal frequency in this group did not cause a compensatory reduction in meal size (Figure 1D). In agreement with (24), olfactory detection of PB did not influence chow consumption in mice (Table S1), probably because of the small amount of food a mouse consumes over a day, and especially during the light phase. PB consumption data are detailed in Table S2.

**Figure 1.**
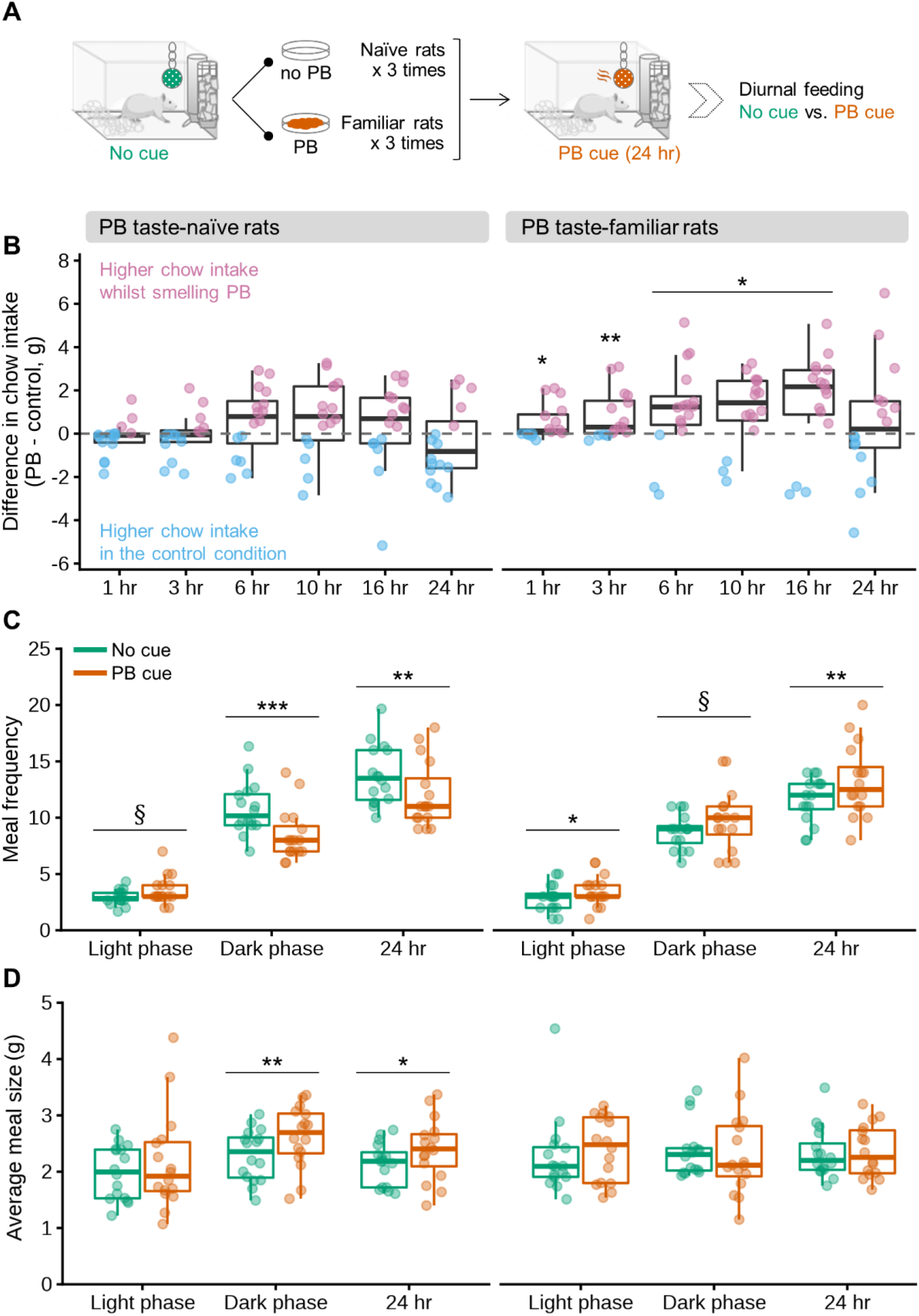
Olfactory detection of PB causes persistent overconsumption of regular chow due to an increase in meal frequency in sated PB taste-familiar rats. **(A)** Schematic of the cross-over experimental design in the automated feeding monitoring cages. **(B)** Difference in chow intake (g) between the PB cue and the control conditions at the 1, 3, 6, 10, 16 and 24 hr time points in PB taste-naïve rats (left, n=16) and PB taste-familiar rats (right, n=16). Animals consuming more chow whilst detecting PB in the environment are represented by pink whiskers, while those who consumed less chow whilst detecting PB in the environment (thus, more chow in the control condition) are represented by blue whiskers. **(C)** Meal frequency and **(D)** average meal size (g) during the light phase, dark phase and total day (24 hr) in PB taste-naïve rats (left, n=16) and PB taste-familiar rats (right, n=16). In all cases, the thick line corresponds to the median, boxes show first and third quartiles and whiskers represent minimum and maximum values. Symbols indicate differences between the control and the PB cue conditions at *p* > 0.05 (*), *p* > 0.01 (**), *p* > 0.001 (***) or 0.05 ≤ *p* < 0.1 (§). PB; peanut butter.

### Olfactory detection of PB prompts food-seeking in sated PB taste-familiar rats

Besides acoustic measurements in the home cage, which were higher in the PB taste-familiar than in the PB taste-naïve group (Table S3), we also explored the incentive salience of PB in a risky environment in the same cohorts of PB taste-naïve and PB taste-familiar rats (Figure 2A). Baiting the center of the arena with PB odor did not evoke food-seeking in PB taste-naïve rats (Figure 2B–C). The duration of the first contact with the perforated ball in the cue-setting did, on the contrary, increase significantly by 55% in rats that were familiar with PB taste (Figure 2B). Likewise, PB taste-familiar rats spent 93% more time exploring the bait (in comparison with the time spent exploring the empty perforated ball) during the entire 10-min test despite the aversive environment (Figure 2C). We did not observe any differences in latency to approach the set-up for the first time for either group (Figure 2D), suggesting that the anxiety-like response was similar among groups (20).

**Figure 2.**
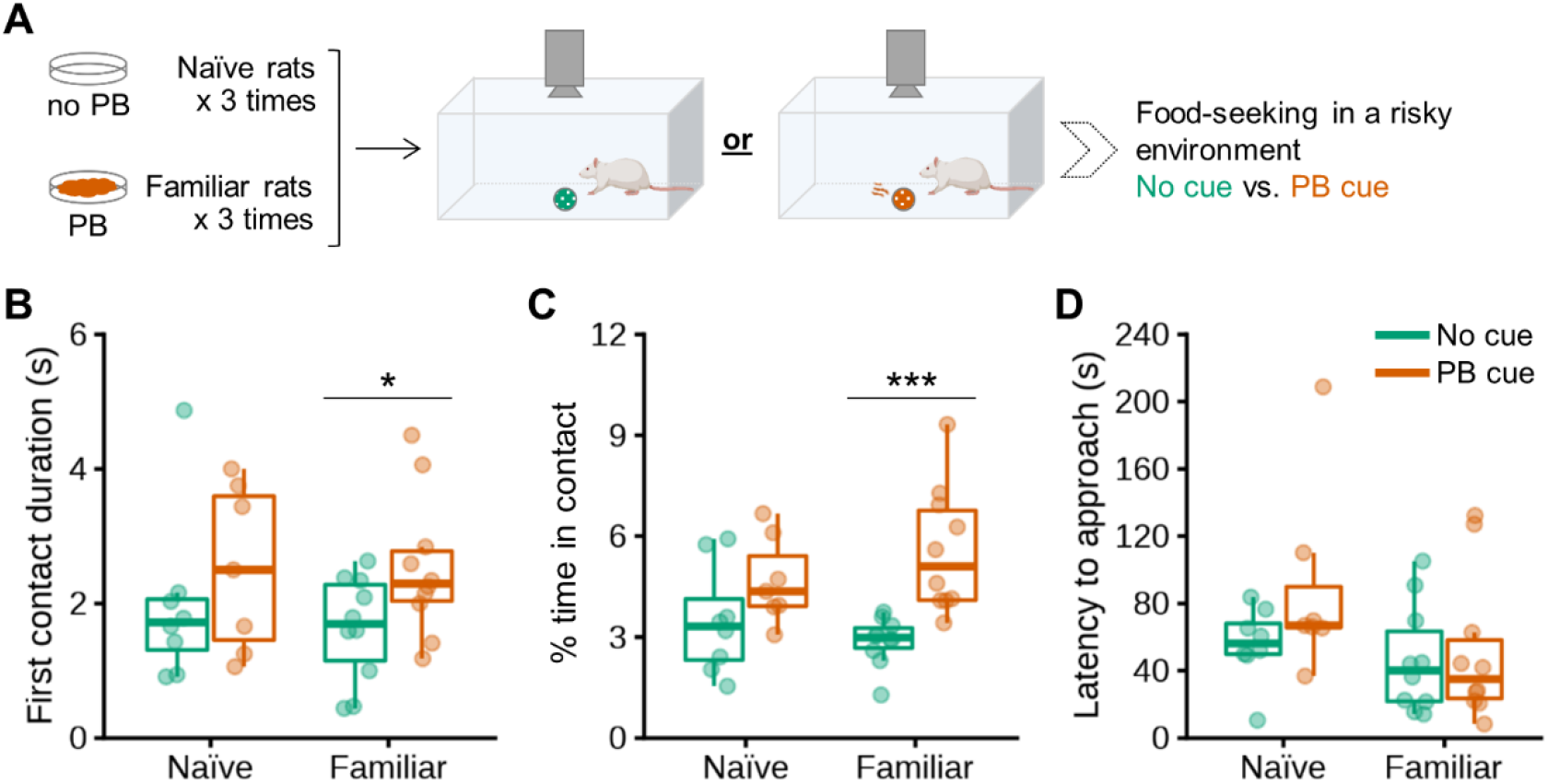
Olfactory detection of PB stimulates food-seeking in a risky environment in sated PB taste-familiar rats. **(A)** Schematic of the experimental design in the PB-baited open field. Effects of baiting the center of the arena with PB on **(B)** first contact duration (s), **(C)** % of time in contact with the perforated ball, as well as **(D)** latency to approach the perforated ball in PB taste-naïve rats (n=15: No cue, n=8; PB cue, n=7) and PB taste-familiar rats (n=20: No cue, n=10; PB cue, n=10). In all cases, the thick line corresponds to the median, boxes show first and third quartiles and whiskers represent minimum and maximum values. Symbols indicate differences between the control and the PB cue conditions at *p* > 0.05 (*) or *p* > 0.001 (***). PB; peanut butter.

### Olfactory detection of PB activates cells in the arcuate nucleus in sated PB taste-familiar rats

To explore the neural substrates engaged in the orexigenic and food-seeking responses to the olfactory PB cue, we first examined whether it activated cells in the Arc. Sated PB taste-naïve and PB taste-familiar rats were exposed to a cue-enriched environment or to a non-enriched environment for 20 min (Figure 3A). Consistent with the food intake data, the olfactory PB cue induced Arc Fos expression only in PB taste-familiar rats (148% more cells compared to the control condition) (Figure 3B–C). Unexpectedly, we were unable to detect more than a few scattered Fos+ cells in any other hypothalamic or brain areas. The only exceptions were the piriform and cingulate cortex, the olfactory tubercle and the medial amygdaloid nucleus, all areas being important relays for olfactory information (25, 26). Fos was detected in these regions in all groups exposed to the olfactory cue, independently of whether they were familiar with PB taste or not (data not shown).

**Figure 3.**
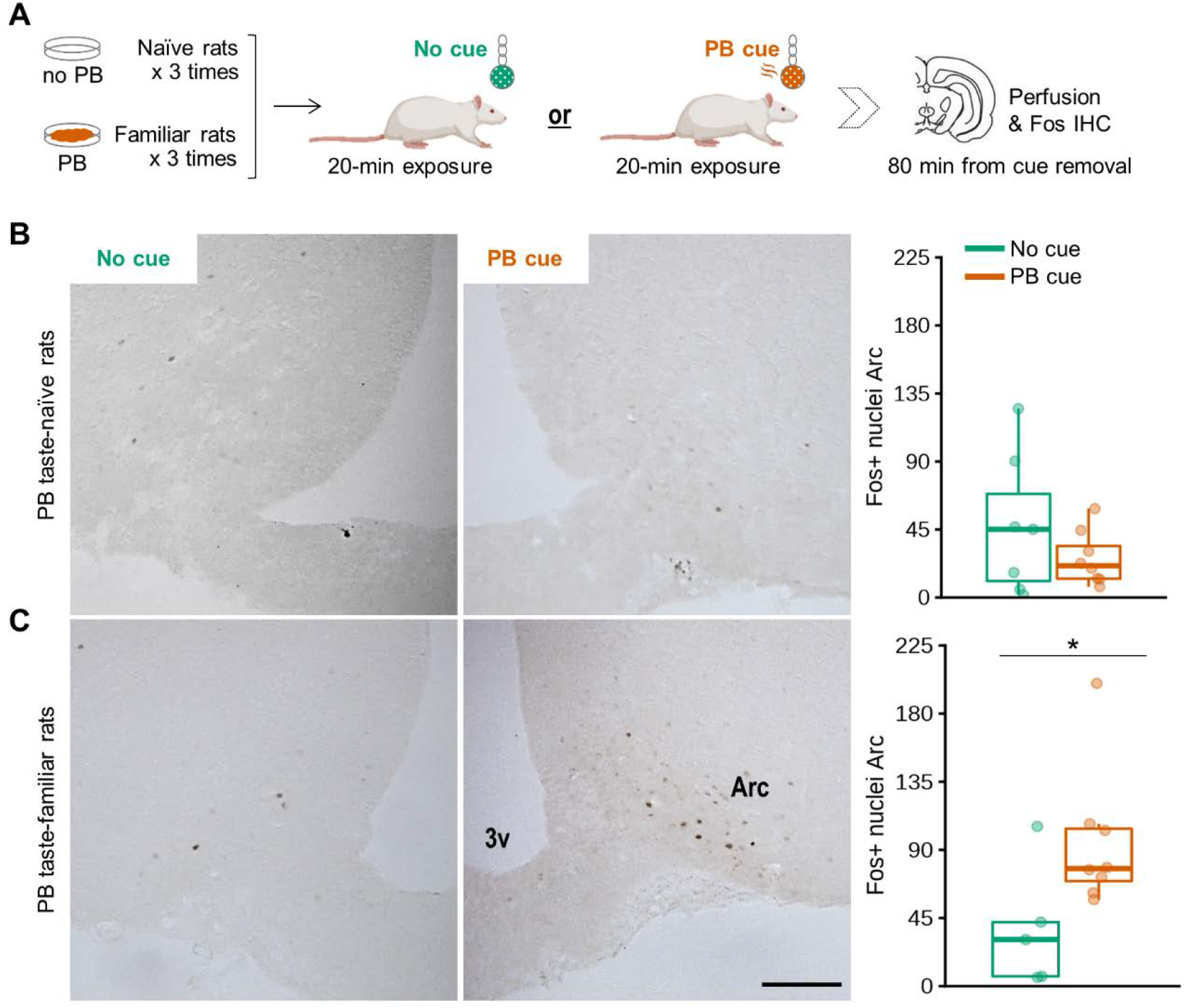
Olfactory detection of PB activates cells in the Arc in sated PB taste-familiar rats. **(A)** Schematic of the experimental design in the home cage. Representative images showing effects of exposure to a 20-min olfactory PB cue to increase the number of cells detected that express Fos protein in the Arc, together with the corresponding unilateral manual counting of Fos+ nuclei in **(B)** PB taste-naïve rats (n=15: No cue, n=7; PB cue, n=8) and **(C)** PB taste-familiar rats (n=13: No cue, n=5; PB cue, n=8). Arc, arcuate nucleus; 3v, third ventricle. Bregma: −2.40 mm; Scale bar = 200 μm (applies to all 4 images). In all cases, the thick line corresponds to the median, boxes show first and third quartiles and whiskers represent minimum and maximum values. Symbols indicate differences between the control and the PB cue conditions at *p* > 0.05 (*). PB; peanut butter.

### Intermingled and overlapping neuronal populations in the Arc are activated upon olfactory detection of PB in sated rats

We then sought to obtain a molecular census of the Fos+ cells in the Arc responding to the olfactory PB cue. The RNAscope study demonstrates that olfactory detection of PB by sated rats does not merely activate a single Arc neuronal population, but rather intermingled population of cells. Of the distinct types of neurons analyzed, GHSR-expressing cells were found to be the most prominent cluster: 35.7 ± 2% of Fos+ cells expressed GHSR (averaged from all the data of the three assays [39.1, 34.9 and 32.1%]; Figures 4C, 5C and 6C). Additionally, as many as 20.5 ± 0.9% of Fos+ cells co-expressed AgRP (Figure 4B), almost a quarter (23.5 ± 1.9%) were POMC+ (Figure 5B), while only 12.6 ± 1.0% were dopaminergic (i.e., expressed *Th* mRNA) (Figure 6B). The proportion of GHSR, AgRP, POMC and TH+ cells that also expressed Fos was 24.8 ± 3.0 (averaged from all the data of the three assays for GHSR [31.6, 23.4 and 16.8%]), 24.2 ± 2.3, 36.1 ± 4.4 and 24.2 ± 6.5%, respectively (Figures 4B, 5B and 6B). We also report additional data about co-localization of GHSR with AgRP, POMC and TH (Figures 4D, 5D, 6D), as well as triple co-expressions (Figures 4F, 5F, 6F) that are described in Supplemental Results.

**Figure 4.**
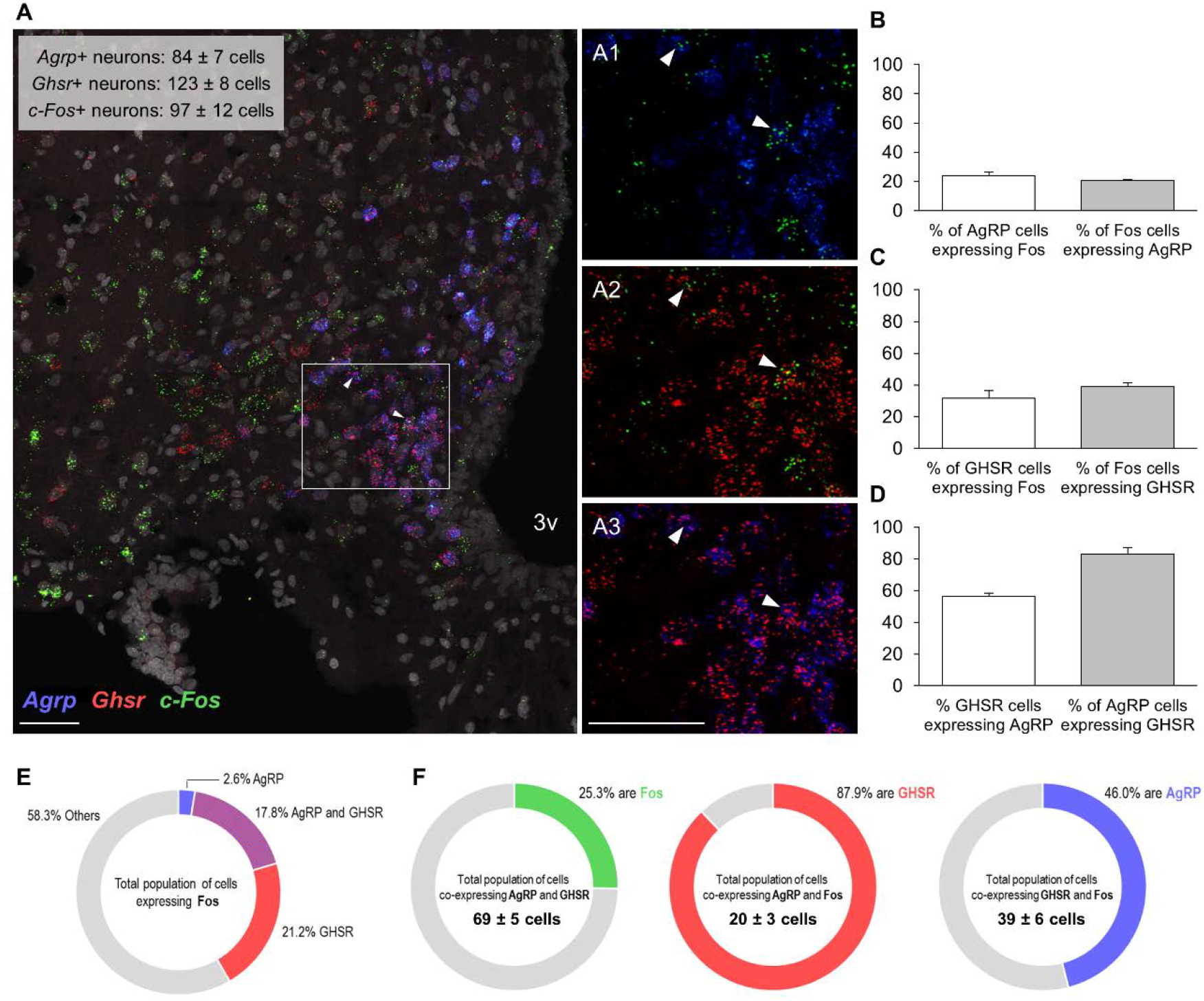
Co-localization of mRNAs for Fos protein, GHSR and AgRP in the Arc of sated PB taste-familiar rats exposed to the olfactory PB cue. **(A)** Representative confocal images of triple RNAscope *in situ* hybridization for *c-Fos* (green), *Ghsr* (red) and *Agrp* (blue) in an Arc-containing section of a PB taste-familiar rat exposed to the olfactory PB cue. Area in the white rectangle is shown enlarged in the small panels on the right **(A1, A2, A3)**. Co-localization of mRNAs for **(A1)** Fos protein and AgRP, **(A2)** Fos protein and GHSR, and **(A3)** GHSR and AgRP. White arrows indicate triple positive cells. The graphs depict **(B)** % of AgRP cells that are Fos+, and the % of Fos cells that are AgRP+; **(C)** % of GHSR cells that are Fos+, and the % of Fos cells that are GHSR+; **(D)** % of GHSR cells that are AgRP+, and the % of AgRP cells that are GHSR+. **(E)** Overview of the molecular identities of Fos+ cells. **(F)** Triple co-localization data. 3v, third ventricle. Bregma: −2.64 mm; Scale bar = 50 μm (applies to all 4 images). 2-4 hemisections per mouse were quantified (n=4).

**Figure 5.**
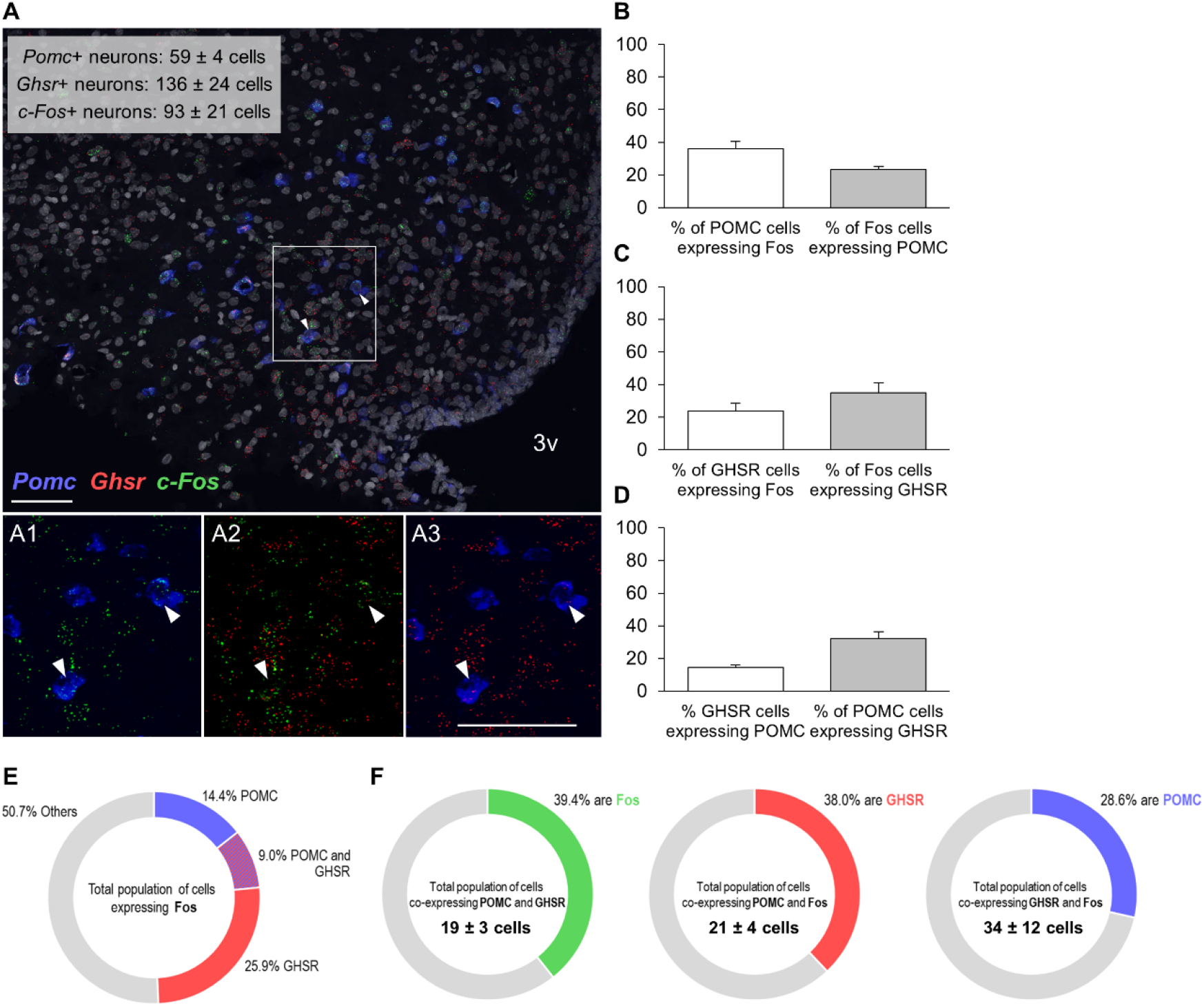
Co-localization of mRNAs for Fos protein, GHSR and POMC in the Arc of sated PB taste-familiar rats exposed to the olfactory PB cue. **(A)** Representative confocal images of triple RNAscope *in situ* hybridization for Fos (green), GHSR (red) and POMC (blue) in an Arc-containing section of a PB taste-familiar rat exposed to the olfactory PB cue. Area in the white rectangle is shown enlarged in the small panels on the bottom **(A1, A2, A3)**. Co-localization of mRNAs for **(A1)** Fos protein and POMC, **(A2)** Fos protein and GHSR, and **(A3)** GHSR and POMC. White arrows indicate triple positive cells. The graphs depict **(B)** % of POMC cells that are Fos+, and the % of Fos cells that are POMC+; **(C)** % of GHSR cells that are Fos+, and the % of Fos cells that are GHSR+; **(D)** % of GHSR cells that are POMC+, and the % of POMC cells that are GHSR+. **(E)** Overview of the molecular identities of Fos+ cells. **(F)** Triple co-localization data. 3v, third ventricle. Bregma: −3.48 mm; Scale bar = 50 μm (applies to all 4 images). 3-4 hemisections per mouse were quantified (n=3).

**Figure 6.**
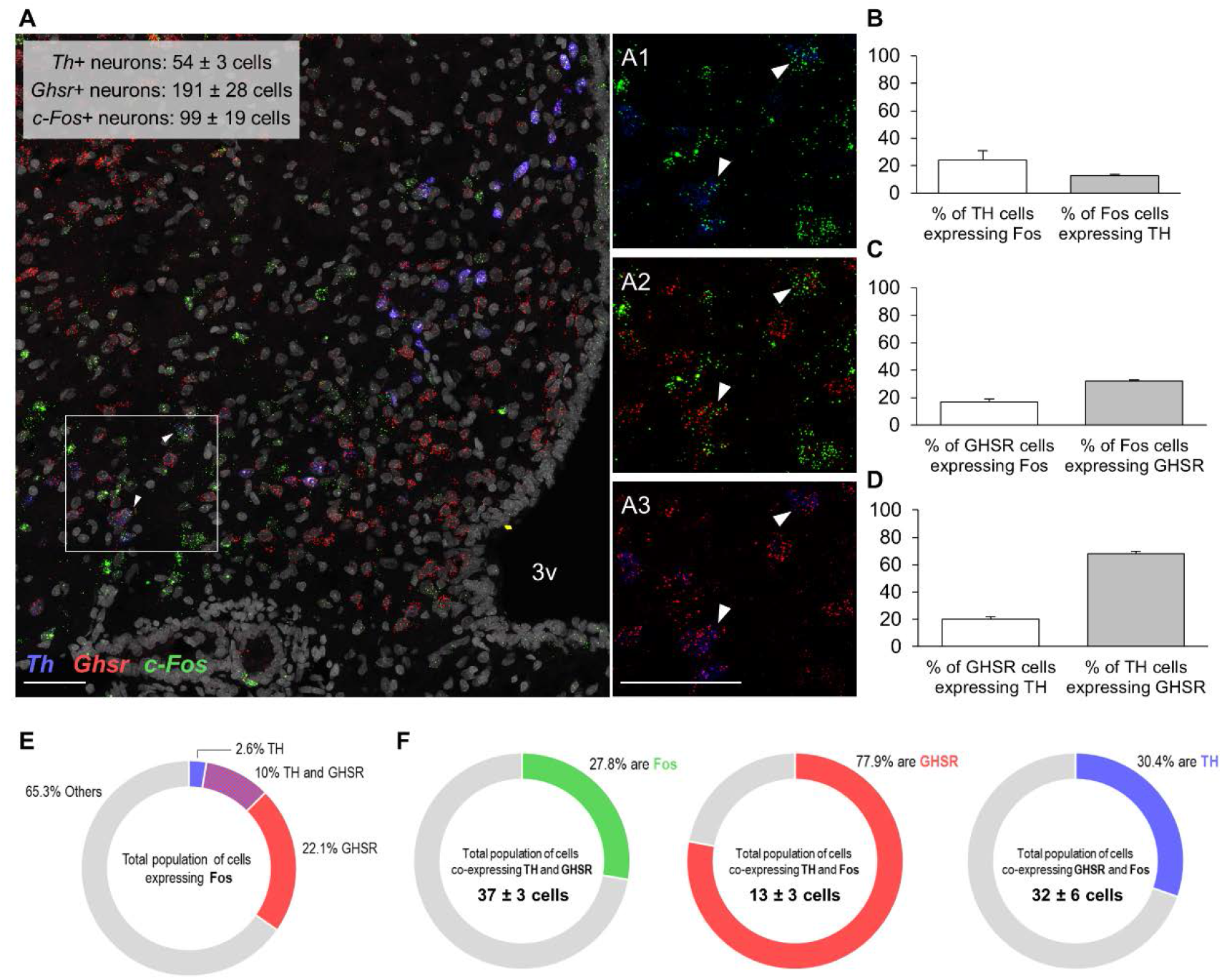
Co-localization of mRNAs for Fos protein, GHSR and TH in the Arc of sated PB taste-familiar rats exposed to the olfactory PB cue. **(A)** Representative confocal images of triple RNAscope *in situ* hybridization for Fos (green), GHSR (red) and TH (blue) in an Arc-containing section of a PB taste-familiar rat exposed to the olfactory PB cue. Area in the white rectangle is shown enlarged in the small panels on the right **(A1, A2, A3)**. Co-localization of mRNAs for **(A1)** Fos protein and TH, **(A2)** Fos protein and GHSR, and **(A3)** GHSR and TH. White arrows indicate triple positive cells. The graphs depict **(B)** % of TH cells that are Fos+, and the % of Fos cells that are TH+; **(C)** % of GHSR cells that are Fos+, and the % of Fos cells that are GHSR+; **(D)** % of GHSR cells that are TH+, and the % of TH cells that are GHSR+. **(E)** Overview of the molecular identities of Fos+ cells. **(F)** Triple co-localization data. 3v, third ventricle. Bregma: −2.64 mm; Scale bar = 50 μm (applies to all 4 images). 3 hemisections per mouse were quantified (n=3).

### Olfactory detection of PB triggers release of active ghrelin in sated PB taste-familiar rats

After reproducing the effects of exposure to PB smell on food intake in a new cohort of PB taste-familiar sated rats (Figure 7A), we also measured their plasma levels of active (acylated) ghrelin, which were elevated by 33% upon olfactory detection of PB (compared to controls; Figure 7B).

**Figure 7.**
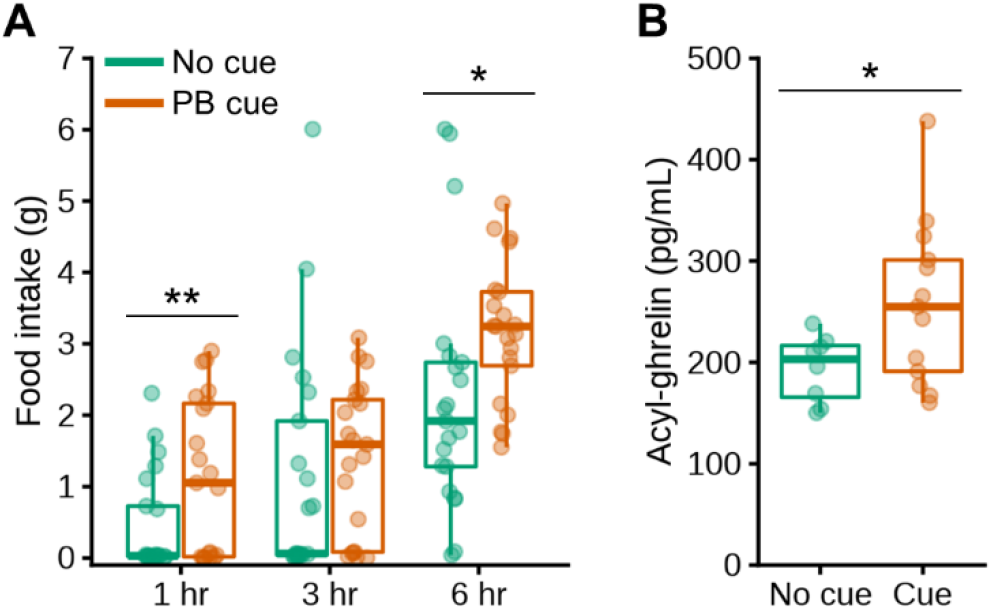
Olfactory detection of PB triggers release of active ghrelin in sated PB taste-familiar rats. **(A)** Confirmation of the cue-induced overeating of chow in a new cohort of PB taste-familiar rats subjected to a cross-over design (n=21). Manual food intake (g) measurements after 1, 3 and 6 hr post-cue introduction in the home cage. The same time points were used for the control condition on the day before, in which spontaneous food intake was measured in the absence of the PB cue following introduction of an empty tube in the perforated ball. **(B)** Levels of acyl-ghrelin (also known as active ghrelin, pg/mL) under sated conditions after 1 hr of exposure to the olfactory PB cue (n=21: No cue, n=8; PB cue, n=13). In all cases, the thick line corresponds to the median, boxes show first and third quartiles and whiskers represent minimum and maximum values. Symbols indicate differences between the control and the PB cue conditions at *p* > 0.05 (*) or *p* > 0.01 (**). PB; peanut butter.

## Discussion

We show that olfactory detection of a familiar palatable food affects diurnal feeding patterns to cause sustained overconsumption of the only readily available food (chow) in sated rats. In line with the orexigenic response observed, detecting PB stimulates food-seeking and triggers release of ghrelin in sated rats. Using Fos mapping, we identify the Arc as a key neural substrate activated by olfactory food cues and provide evidence that these recruit intermingled populations of cells embedded within the feeding circuitry, even when neither chow nor the cued PB are available for consumption.

It is widely accepted that cues recalling food, such as its sight or smell, prompt food-seeking and initiate feeding despite satiation in humans. Satiated subjects, when exposed to a virtual food cue-rich environment, self-report increased hunger and an increased desire to eat, leading to heightened motivation for food and increased caloric consumption (2). The pioneering works of Weingarten first and Petrovich more recently have also demonstrated that both discrete and contextual cues potentiate feeding in rodents through classical Pavlovian conditioning (3–9). In these settings, however, there exists a phase of associative learning that takes place in behavioral chambers and the ultimate feeding response is specific to the signaled food. Our results show that olfactory detection of a palatable food in the home cage is powerful enough to cause chow to be overconsumed, a less preferred food that lacks incentive value when not hungry. Boggiano and colleagues (27) previously demonstrated that context cues concomitantly present whilst consuming a palatable food (e.g., a bedding that differs from the animal’s habitual cage) become drivers to promote overeating of chow when sated rats are introduced in this food-paired context. The perforated ball in our study was already present in the home cage prior to single-housing, and all the baseline food intake measurements took place after the PB tastings, in such a way that any contextual cues (i.e., sudden presence of a new object in the cage or associative contextualization of the home cage with PB intake) could not have influenced the results. In a nutshell, our behavioral approach (18) allowed us not only to investigate long-term feeding outcomes in response to food-linked olfactory cues with small disturbance to the animals, but also to selectively isolate the cue-dependent effects from the associative learning context, which *per se* influences food intake (27).

The cue-driven orexigenic response was already detected after 1 hr, suggesting that it develops rapidly upon cue presentation, but spanned over 16 hr, indicating that it persists even long after PB smell is detected for the first time. Questions that inevitably arises are what drives rats in our model to respond to the PB cue by overeating chow, and why is this response sustained over time? Petrovich and colleagues, in their work, discussed the fact that the cue-driven orexigenic response underlies a specific motivation for the signaled food, akin to craving (5). Our results suggest that sensory detection of PB in the environment drives a general desire to eat, similar to that expected from experiencing hunger. The motivation for initiating a meal certainly included an early stage of specific craving for PB, which conceivably preceded the assimilation by the rats that the olfactory cue did not signal its access. Without further expectations of obtaining PB, the rats’ physiology may have worked towards fulfilling this new set point by fostering the rats to overconsume the available chow. The fact that chow intake was not greater after 24 hr, however, suggests that the animals eventually compensate for their earlier cue-linked overconsumption, and that energy homeostasis prevails, as already postulated decades ago by Weingarten (9). Importantly, however, this substantial intake of chow, albeit transient, was not compensated for following 24 hr cue removal, because daily chow intake post-cue was similar to both the baseline and the cue settings.

This temporary overconsumption appears to reflect especially an increased meal frequency that was inadequately compensated for. Such a higher frequency of meals likely underscores an effect of olfactory food-linked cues in selectively activating appetitive systems regulating meal initiation within hypothalamic areas, whereas a failure to decrease meal size would indicate engagement of hindbrain pathways involved in satiation and meal termination (28). Together with our finding that circulating active ghrelin levels are increased 1 hr after the presentation of the cue, and consistent with a role of endogenous ghrelin as a physiological meal initiator (29, 30), these data would substantiate an engagement of the ghrelin system in driving, at least acutely, the appetitive and orexigenic components of the behavioral response triggered by the cue. Corroborating this idea, evidence from human studies link an enhanced response to primary food-linked cues (i.e., food pictures) with heightened circulating ghrelin (31). Our data, therefore, support the notion that there exists a feed-forward mechanism whereby olfactory food cues engage the ghrelin system by enhancing ghrelin release, which eventually heightens food-cue sensitivity (arguably by boosting olfactory acuity (17)), food-seeking and feeding, behaviors that are well-known to be driven by this hormone (20, 22, 32). The level at which this feed-forward mechanism occurs is unclear. It is tempting to speculate that it might be fueled by the direct action of ghrelin via its receptor on olfactory processing centers, as well as through its positive and inhibitory actions on AgRP and POMC neurons, respectively (33). Yet another possibility is that olfactory inputs, through olfactory circuits, communicate directly with central neurons to regulate both food-seeking and ingestion.

Due to the pronounced food-seeking for the cued PB, we investigated the potential of such olfactory cues on influencing risk-taking in a PB-baited open field test. It seems clear that this appetitive state directs the animal to, and motivates initial contact with, the cued food: the PB bait in the center of the field shifted the individual’s behavioral choice towards a more risk-taking phenotype, because animals spent twice as much time exploring the perforated ball despite the anxiogenic environment. Consistently, both fasting and exogenous ghrelin have been found to reset the perceived pressure on food supply, thus encouraging sated mice to forgo the fear of an open field over the chance of getting food (20).

The ensemble of responses to the cued PB suggest that olfactory food cues engage different pathways involved in motivation, feeding and energy homeostasis. Consistent with this, we found that a 20-min exposure to PB smell activated cells in the Arc in sated rats, similar to that observed for other orexigenic stimuli such as fasting (34) and ghrelin administration (21). We were surprised by the fact that the olfactory PB cue did not induce Fos protein expression in any other hypothalamic or reward-linked areas. Indeed, it would be expected from exposure to cues evoking palatable foods that they increase responsiveness of the reward circuitry to promote the seeking and the subsequent cue (specific)-potentiated feeding (35). Yet the lack of Fos expression may be weak evidence for lack of a change, and we cannot exclude the involvement of brain areas where neuronal activation is not coupled to a Fos response.

Interestingly, GHSR-containing cells were found to be the more prominent cluster from all the cell types activated upon exposure to PB smell. Did the olfactory cue activate GHSR-containing cells or was it a downstream consequence of cue-induced ghrelin release? Based on available information, it would be difficult to estimate the extent to which cue-induced ghrelin may have directly contributed to mediate the effects of the olfactory cue as a part of our hypothesized feed-forward mechanism.

Whereas AgRP and POMC neurons have been so far considered to be sensors of energy requirements, the work of Chen and colleagues (15) clearly shows that their activity can be rapidly modulated by the chemosensory detection of food. These observations pave the way towards a new concept in which food-seeking would be a higher order consequence of detecting food in the environment, rather than a downstream consequence of sensing an energy deficit. In line with this, we show that olfactory detection of palatable food in the environment activates both AgRP and POMC neurons to a similar extent. Interestingly, the ghrelin system seems to be an important liaison between *hunger* and responsiveness to olfactory cues: the majority of the recruited AgRP cells also expressed GHSR, while a substantial proportion of POMC did so too. It is worth recalling, at this juncture, that although Fos expression indicates a change in neuronal activity, it does not always reflect increased electrical activity (36). Then, as introduced above, the triple co-expression results might likewise reflect a complex crosstalk between the ghrelin system and external olfactory food-evoking signals, since they might underlie a direct effect of cue-induced ghrelin on AgRP and POMC cells.

Although previous work has shown that dopaminergic cells in the Arc respond to fasting and peripheral ghrelin, and thus probably stand as a novel actor regulating energy homeostasis and food intake (23), our results suggest that these neuronal population might not be of critical relevance for integrating food-linked sensory stimuli in the olfactory modality.

It is worth emphasizing that only rats conditioned to PB taste displayed the cue-evoked motivational and feeding responses. Merely challenging PB taste-naïve rats with PB smell was insufficient to initiate interest for the hidden food or boost chow intake, suggesting that the odor only gains value when the animals are familiar with the taste and ingestion. Although rodents might sense the intrinsic hedonic value of nutritive stimuli (e.g., fat or sugar), this might be not enough to condition a taste preference, and thus post-ingestive actions of nutrients must take place (37), underpinning the role of learning and associative processes in cue-potentiated feeding (3–9).

Collectively, our data endorse a role of ubiquitous olfactory cues evoking palatable foods not only in driving food-seeking (most probably directed towards the signaled food) and release of ghrelin, which appears to be a prominent and versatile actor, but also ultimate feeding, which develops towards the only food available, even if it is non-salient. Our finding that populations of Arc cells embedded within the energy balance – hunger neurocircuitry are engaged by odor food cues even in the sated state, suggest a role for these neurons in food cue-potentiated feeding behaviors. We show that besides AgRP and POMC neurons, which are similarly activated, GHSR-containing and dopaminergic cells are also recruited by the external olfactory food-linked stimulus. One thing is certain: food cues in our society are unavoidable and any failure to adjust for even small increases in intake might lead to caloric surfeit. However, what if the *only* food available appeared to be a healthier choice, as it was the case herein? These results likewise evince the importance of health nudge interventions conducted by governments worldwide, which try to steer people into healthier lifestyles and food choices (38).

## Acknowledgements and Disclosures

This work was supported by the Swedish Research Council for Medicine and Health (2018-02588 to RAHA and 2019-01051 to SLD), the Novo Nordisk Foundation (NNF17OC0027206 and NNF19OC0056694 to SLD), Hjärnfonden (FO2017-0180; FO2018-0262; FO2019-0086 to SLD) and the Swedish state under the agreement between the Swedish Government and the county councils in the ALF agreement (ALFGBG-723681 to SLD). FP-S conceived, designed and led the studies in discussion with SLD and RAHA. Behavioral experiments and ghrelin assessment were conducted by FP-S, assisted by IS and MVL. IS performed immunohistochemistry and imaging. MVL performed the RNAscope experiment and imaging. FP-S did the counting, analyzed, and plotted data. PS-N did the graphics in R and assisted in data analysis. FP-S wrote the manuscript, with inputs from SLD. All authors critically contributed to and ultimately accepted the last version for publication. We thank Dr Tina Bake and Dr Erik Schéle for assistance with feeding experiments in the automated cages and with technical aspects of the animal work, respectively, as well as ECNP for supporting the Nutrition Network. The authors report to have no biomedical financial interest or other potential conflicts of interest.

## References

1. Holland PC, Petrovich GD (2005): A neural systems analysis of the potentiation of feeding by conditioned stimuli. Physiology & behavior. 86:747–761.

2. Joyner MA, Kim S, Gearhardt AN (2017): Investigating an Incentive-Sensitization Model of Eating Behavior: Impact of a Simulated Fast-Food Laboratory. Clinical Psychological Science. 5:1014–1026.

3. Petrovich GD, Holland PC, Gallagher M (2005): Amygdalar and prefrontal pathways to the lateral hypothalamus are activated by a learned cue that stimulates eating. The Journal of neuroscience : the official journal of the Society for Neuroscience. 25:8295–8302.

4. Petrovich GD, Ross CA, Gallagher M, Holland PC (2007): Learned contextual cue potentiates eating in rats. Physiology & behavior. 90:362–367.

5. Petrovich GD, Ross CA, Holland PC, Gallagher M (2007): Medial prefrontal cortex is necessary for an appetitive contextual conditioned stimulus to promote eating in sated rats. The Journal of neuroscience : the official journal of the Society for Neuroscience. 27:6436–6441.

6. Petrovich GD, Hobin MP, Reppucci CJ (2012): Selective Fos induction in hypothalamic orexin/hypocretin, but not melanin-concentrating hormone neurons, by a learned food-cue that stimulates feeding in sated rats. Neuroscience. 224:70–80.

7. Reppucci CJ, Petrovich GD (2012): Learned food-cue stimulates persistent feeding in sated rats. Appetite. 59:437–447.

8. Weingarten HP (1983): Conditioned cues elicit feeding in sated rats: a role for learning in meal initiation. Science (New York, NY). 220:431–433.

9. Weingarten HP (1984): Meal initiation controlled by learned cues: basic behavioral properties. Appetite. 5:147–158.

10. Fine LG, Riera CE (2019): Sense of Smell as the Central Driver of Pavlovian Appetite Behavior in Mammals. Frontiers in physiology. 10:1151.

11. Palouzier-Paulignan B, Lacroix MC, Aimé P, Baly C, Caillol M, Congar P, et al. (2012): Olfaction under metabolic influences. Chemical senses. 37:769–797.

12. Gascuel J, Lemoine A, Rigault C, Datiche F, Benani A, Penicaud L, et al. (2012): Hypothalamus-olfactory system crosstalk: orexin a immunostaining in mice. Frontiers in neuroanatomy. 6:44.

13. Russo C, Russo A, Pellitteri R, Stanzani S (2018): Ghrelin-containing neurons in the olfactory bulb send collateralized projections into medial amygdaloid and arcuate hypothalamic nuclei: neuroanatomical study. Experimental brain research. 236:2223–2229.

14. Schneider NY, Chaudy S, Epstein AL, Viollet C, Benani A, Pénicaud L, et al. (2020): Centrifugal projections to the main olfactory bulb revealed by transsynaptic retrograde tracing in mice. The Journal of comparative neurology. 528:1805–1819.

15. Chen Y, Lin YC, Kuo TW, Knight ZA (2015): Sensory detection of food rapidly modulates arcuate feeding circuits. Cell. 160:829–841.

16. Mani BK, Osborne-Lawrence S, Mequinion M, Lawrence S, Gautron L, Andrews ZB, et al. (2017): The role of ghrelin-responsive mediobasal hypothalamic neurons in mediating feeding responses to fasting. Molecular metabolism. 6:882–896.

17. Tong J, Mannea E, Aimé P, Pfluger PT, Yi CX, Castaneda TR, et al. (2011): Ghrelin enhances olfactory sensitivity and exploratory sniffing in rodents and humans. The Journal of neuroscience : the official journal of the Society for Neuroscience. 31:5841–5846.

18. Stoltenborg I, Peris-Sampedro F, Le May MV, Bake T, Schele E, Adan RA, et al. (2020): Primary food cues engage pathways involved in over-eating and reward-seeking in rats. European Neuropsychopharmacology, 28 January, 2020 ed, pp S 36–S 37.

19. Crossley E, Biggs T, Brown P, Singh T (2021): The Accuracy of iPhone Applications to Monitor Environmental Noise Levels. The Laryngoscope. 131:E59–e62.

20. Lockie SH, McAuley CV, Rawlinson S, Guiney N, Andrews ZB (2017): Food Seeking in a Risky Environment: A Method for Evaluating Risk and Reward Value in Food Seeking and Consumption in Mice. Frontiers in neuroscience. 11:24.

21. Peris-Sampedro F, Stoltenborg I, Le May MV, Zigman JM, Adan RAH, Dickson SL (2021): Genetic deletion of the ghrelin receptor (GHSR) impairs growth and blunts endocrine response to fasting in Ghsr-IRES-Cre mice. Molecular metabolism. 51:101223.

22. Wren AM, Small CJ, Abbott CR, Dhillo WS, Seal LJ, Cohen MA, et al. (2001): Ghrelin causes hyperphagia and obesity in rats. Diabetes. 50:2540–2547.

23. Zhang X, van den Pol AN (2016): Hypothalamic arcuate nucleus tyrosine hydroxylase neurons play orexigenic role in energy homeostasis. Nature neuroscience. 19:1341–1347.

24. Boone MH, Liang-Guallpa J, Krashes MJ (2021): Examining the role of olfaction in dietary choice. Cell reports. 34:108755.

25. Ciorba A, Hatzopoulos S, Cogliandolo C, Bianchini C, Renna M, Pelucchi S, et al. (2020): Functional Magnetic Resonance Imaging in the Olfactory Perception of the Same Stimuli. Life (Basel, Switzerland). 11.

26. Wilson DA, East BS (2021): Good scents: A short road from olfaction to satisfaction. Current biology : CB. 31:R374–r376.

27. Boggiano MM, Dorsey JR, Thomas JM, Murdaugh DL (2009): The Pavlovian power of palatable food: lessons for weight-loss adherence from a new rodent model of cue-induced overeating. International journal of obesity (2005). 33:693–701.

28. Grill HJ (2006): Distributed neural control of energy balance: contributions from hindbrain and hypothalamus. Obesity (Silver Spring, Md). 14 Suppl 5:216s–221s.

29. Cummings DE (2006): Ghrelin and the short-and long-term regulation of appetite and body weight. Physiology & behavior. 89:71–84.

30. Faulconbridge LF, Cummings DE, Kaplan JM, Grill HJ (2003): Hyperphagic effects of brainstem ghrelin administration. Diabetes. 52:2260–2265.

31. Schüssler P, Kluge M, Yassouridis A, Dresler M, Uhr M, Steiger A (2012): Ghrelin levels increase after pictures showing food. Obesity (Silver Spring, Md). 20:1212–1217.

32. Merkestein M, Brans MA, Luijendijk MC, de Jong JW, Egecioglu E, Dickson SL, et al. (2012): Ghrelin mediates anticipation to a palatable meal in rats. Obesity (Silver Spring, Md). 20:963–971.

33. Cowley MA, Smith RG, Diano S, Tschop M, Pronchuk N, Grove KL, et al. (2003): The distribution and mechanism of action of ghrelin in the CNS demonstrates a novel hypothalamic circuit regulating energy homeostasis. Neuron. 37:649–661.

34. Riediger T, Bothe C, Becskei C, Lutz TA (2004): Peptide YY directly inhibits ghrelin-activated neurons of the arcuate nucleus and reverses fasting-induced c-Fos expression. Neuroendocrinology. 79:317–326.

35. Volkow ND, Wang GJ, Tomasi D, Baler RD (2013): Obesity and addiction: neurobiological overlaps. Obesity reviews : an official journal of the International Association for the Study of Obesity. 14:2–18.

36. Johnstone LE, Fong TM, Leng G (2006): Neuronal activation in the hypothalamus and brainstem during feeding in rats. Cell metabolism. 4:313–321.

37. Myers KP, Sclafani A (2006): Development of learned flavor preferences. Developmental psychobiology. 48:380–388.

38. Leng G, Adan RAH, Belot M, Brunstrom JM, de Graaf K, Dickson SL, et al. (2017): The determinants of food choice. The Proceedings of the Nutrition Society. 76:316–327.

