## Supplementary Information for "The orexigenic force of olfactory palatable food cues in sated rats"

**SUPPLEMENTAL INFORMATION**

**Supplementary Methods and Materials**

**Animals**

Male Sprague-Dawley rats (n=88), aged 7/9-week-old and weighing 322 to 406 g upon arrival, were purchased from Charles River (Calco, Italy). Male C57BL/6N mice (n=10), aged 12-week-old and weighing 30.3 ± 1.1 g upon arrival, were also purchased from Charles River. Animals were allowed one-week acclimatization to the facility prior to being single-housed. They had *ad libitum* access to water and chow (2016 Teklad diet, Harlan Laboratories, Cambridgeshire, UK), unless otherwise stated, and were kept on a 12-hr light-dark cycle (lights on: 7 AM) for the duration of the studies. In all cases, a metallic, perforated, opaque tea-strainer ball was suspended from each animal’s cage lid (1) prior to single housing to avoid neophobia (to ensure that the animals were familiar with the set-up before each experiment started) and to prevent it from becoming a non-food related contextual cue (2). During experiments, this device contained the inaccessible palatable food (peanut butter, PB; Skippy, Hormel foods, Austin, MN, USA), delivering an olfactory PB cue to the home environment. The choice of PB was based on pilot studies in which we determined the preference for different palatable foods, but also on its organoleptic properties. To avoid cross-odor contamination, experiments in PB taste-naïve rats were carried out first (and on different days) from those in PB taste-familiar rats. Only males were used because of unknown impacts of the estrus cycle in females on the parameters measured.

**Feeding response and meal patterns in an olfactory PB cue-enriched environment**

An automated feeding monitoring system (TSE LabMaster, Project 4261, TSE Systems, Bad Homburg, Germany) was used to analyze diurnal feeding patterns (measuring cumulative chow intake [g], meal frequency, meal size [g], and ingestion rate [g/min]) in an olfactory PB cue-enriched environment in both PB taste-naïve (n=16) and PB taste-familiar (n=16) rats. This system allows uninterrupted and undisturbed recording of individual meals for each animal. Food hoppers containing regular pelleted chow were suspended on sensors (calibrated prior to starting the experiments) recording food intake to a sensitivity of 1 mg. Rats were transferred into the cages and allowed one-week habituation prior to starting the feeding recordings.

Novel olfactory PB-cue stimulus (PB taste-naïve rats)

First, we generated PB taste-naïve rats (n=16). To keep the familiarization procedure as similar as possible between PB taste-naïve and PB taste-familiar groups, rats were presented with an inedible object (empty food dish) once a day for three alternating days after habituation to the experimental cage. To avoid interval-dependent anticipation, the object was presented at different time points and no identical intervals were repeated. Afterwards, baseline chow intake was calculated based on the daily automatically recorded feeding measurements. Data were averaged from the last two days of measurements and reported as 1-, 3-, 6-, 10-, 16- and 24-hr chow intake. Afterwards, open tubes containing PB were encased in the perforated balls hanging from each of the lids immediately after the light cycle onset and left there for 24 hr. Cumulative chow consumption in the olfactory cue-enriched environment (1, 3, 6, 10, 16 and 24 hr post-cue introduction), as well as within 24 hr following cue removal, was recorded and ultimately compared to that of the baseline period.

Conditioned olfactory PB-cue stimulus (PB taste-familiar rats)

Two weeks later, another cohort of rats (n=16) were exposed to PB tasting. Briefly, they were subjected to the same familiarization protocol as the PB taste-naïve rats, but they were given 1-hr access to consume a limited amount of PB for three alternating days, at varying time intervals, instead of the inedible object. The amount of PB consumed was measured after each tasting session. After baseline food measurements, PB-filled tubes were encased in the perforated balls as described above. Cumulative chow consumption in the olfactory cue-enriched environment (1, 3, 6, 10, 16 and 24 hr post-cue introduction), as well as within 24 hr following cue removal, was recorded and ultimately compared to that of the baseline period.

Meal pattern analysis in the novel and the conditioned settings

Data for meal pattern analysis were collected as binary data every 10 sec. Meal analysis was undertaken using LabMaster software (TSE Systems), whereby all meals occurring during the study period (baseline and 24-hr exposure to the cue-enriched environment in both PB taste-naïve and PB taste-familiar groups) were recorded chronologically to allow the evaluation of single feeding bouts. Meals were defined as an episode of feeding in which at least 0.5 g of chow was removed, with meal termination criterion as the beginning of at least 10-min pause in ingestion (3). Ingestion rate during a meal was calculated by dividing meal size by meal duration. Meal frequency and meal size were summarized over different periods: light phase (12 hr), dark phase (12 hr), and total day (24 hr), and then averaged per rat and group.

Cue-induced feeding response in PB taste-familiar mice

Mice (n=10) were first familiarized with PB taste as described for PB taste-familiar rats. The amount of PB consumed was likewise measured after each tasting session. A tube containing PB was encased in each of the perforated balls and chow intake was measured manually (using scales that had a precision of 1 mg) at 1, 3 and 6 hr post-cue introduction. The same time points were used for the control condition on the day before, in which we measured spontaneous food intake in the absence of the olfactory PB cue following introduction of an empty tube in the perforated ball.

**Food-seeking in familiar and risky environments enriched with an olfactory PB cue**

Acoustic measurements in the home environment

All rats (PB taste-naïve, n=15; PB taste-familiar, n=20) were initially brought to the experimental room 30 min prior to starting the recordings for acclimatization. All the recordings were carried out using the Sound Level Meter app (4) and took place early in the morning (~ 8 AM), to avoid the bustle of the facility, and at the same time for each cohort. After recording the baseline noise (minimum, average and maximum intensities [dB]) following introduction of an empty tube in each of the perforated balls, the latter was replaced with a PB-filled tube and the environmental noise was then recorded (minimum, average and maximum intensities [dB]) in the presence of PB odor. The sound intensity difference (∆ dB) between the baseline and the cue-setting recordings was calculated for the total 15-min period.

PB-baited open field test

Rats from each cohort were then subdivided into the following groups: PB taste-naïve/PB cue (n=7), PB taste-naïve/No cue (n=8), PB taste-familiar/PB cue (n=10) and PB taste-familiar/No cue (n=10). The PB-baited open field test was adapted from a validated protocol (5), and was used to assess risk-taking behavior in food-seeking. The open field boxes consisted of two different bright-white 900 × 900 mm arenas, protected with 300 mm-high walls (Med Associates Inc., St Albans, Vermont, USA). On the day of the experiment, either an empty or a PB-containing perforated ball was placed in the center of the field and secured to the floor. Each rat was initially placed in the left corner of the field and allowed to freely explore the space without interruption for 10 min. Each trial was filmed for later video analysis, which was carried out manually. To ease the analysis, we draw a virtual “cue zone” (the area immediately surrounding the perforated ball), which was kept constant for all the trials. Latency was defined as time to approach the perforated ball after introduction into the arena. The duration of the first contact with the perforated ball, as well as the total time spent in contact with the set-up were measured. The perforated ball was cleaned with ethanol between each trial. The behavioral assessment took place during the light phase (10 AM to 2 PM) and groups were counterbalanced regarding the time of testing.

**Cell activation in the arcuate nucleus upon olfactory detection of PB**

To boost the associative reinforcement, all PB taste-familiar rats were given an extra 1-hr access to consume PB on the day prior to sacrifice. To avoid anticipation, the subgroups subjected to the open field test were swapped for this experiment. The groups were then as follows: PB taste-naïve/PB cue (n=8), PB taste-naïve/No cue (n=7), PB taste-familiar/PB cue (n=8), and PB taste-familiar/No cue (n=5).

Immunohistochemistry and imaging

At 80 min from cue removal, rats were deeply anesthetized with a mixture of Rompun vet. ® (10 mg/kg; Bayer, Leverkusen, Germany) and Ketaminol vet. ® (75 mg/kg; Intervet, Boxmeer, Netherlands), prior to being perfused transcardially with heparinized 0.9 % saline followed by 4 % paraformaldehyde (PFA) in 0.1 M phosphate-buffered saline (PBS), following a group-counterbalanced timing. After harvesting, the brains were stored overnight at 4°C in a 4 % PFA fixative solution containing 15 % sucrose, followed by a minimum of 12-hr incubation in 30% sucrose/0.1 M PBS to ensure cryoprotection. Coronal sections (30-μm thick) containing the Arc were cut using a cryostat and stored in an antifreeze solution (25 % glycerine, 25 % ethylene glycol, 50 % 0.1 M PBS) at -20°C until further processing.

Free-floating sections were processed for the immunohistochemical detection of Fos protein using the 3,3′-diaminobenzidine (DAB)-hydrogen peroxidase method (6). After deactivation of endogenous peroxidases, the sections were rinsed with 0.1 M PBS + 0.3% Triton X-100 prior to being blocked for 1 hr at room temperature in 0.1 M PBS, 3% normal goat serum, 0.25% BSA and 0.3% Triton X-100. Afterwards, they were incubated with an anti-c-Fos rabbit primary antibody (dilution 1:20 000; Ab‐5 (4‐17) Rabbit pAb, PC38; Calbiochem, San Diego, CA, USA) for three nights. The sections were then rinsed and subsequently incubated for 2 hr with a peroxidase goat anti-rabbit immunoglobulin (Ig)G secondary antibody (dilution 1:200; PI1000; Vector Laboratories, Burlingame, CA, USA) and a DAB, nickel, and hydrogen peroxide solution. Brain sections were mounted onto glass slides and coverslipped with ProLong® Diamond Antifade mountant (Thermo Fisher, Waltham, MA, USA).

Unilateral images (5 sections per rat from 2.04 to 3.72 caudal to Bregma) were acquired from rostral to caudal using a DMRB fluorescence microscope (10X/N.A. 0.30; Leica Microsystems, Wetzlar, Germany). The number of Fos+ cells per section was counted manually in ImageJ/Fiji (NIH, Bethesda, MD, USA) using the Cell counter plug-in. The mean number of Fos+ cells per hemisection (and averaged per three blind countings) was calculated, averaged for each brain and ultimately for each experimental group.

**Neurochemical identification of the cells activated by the olfactory PB cue**

Triple fluorescent *in situ* hybridization using RNAscope® (6) was performed to study the potential co-expression of Fos (*c-Fos* probed) with ghrelin receptor (GHSR)- (*Ghsr* probed), agouti-related protein (AgRP)- (*Agrp* probed), Pro-opiomelanocortin (POMC)- (*Pomc* probed) and dopamine (*Tyrosine hydroxylase* *[Th]* probed)-containing cells in the Arc of PB taste-familiar rats (n=3-4) upon a 20-min exposure to PB odor. To this end, three independent assays were run: (i) *c-Fos*, *Ghsr* and *Agrp*; (ii) *c-Fos*, *Ghsr* and *Pomc*; and (iii) *c-Fos*, *Ghsr* and *Th*. This also allowed us to study the co-localization of GHSR with AgRP, POMC and TH. Rats were deeply anesthetized with the mixture of Rompun vet. ® and Ketaminol vet. ® and perfused transcardially, as previously described. After harvesting, the brains were stored overnight at 4°C in a 4 % PFA fixative solution, and then kept in 0.1 M PBS containing 25 % sucrose at 4°C until cryosection. Coronal sections containing the Arc (14 μm-thick, every 6th section collected to provide six adjacent series) were cut using a cryostat and stored in an antifreeze solution (25 % glycerine, 25 % ethylene glycol, 50 % 0.1 M autoclaved PBS) at -20°C until further processing.

All reagents were purchased from Advanced Cell Diagnostics (ACD, Hayward, CA, United States) if not stated otherwise. The *c-Fos* probe (#403591-C3) contained 20 oligonucleotide pairs and targeted region 473-1497 (Acc. No. NM_022197.2) of the *c-Fos* transcript. The *Ghsr* probe (#480031) contained 14 oligonucleotide pairs and targeted region 2-742 (Acc. No. NM_032075.3) of the *Ghsr* transcript. The *Agrp* probe (#316171-C2) contained 13 oligonucleotide pairs and targeted region 14-613 (Acc. No. NM_033650.1) of the *Agrp* transcript. The *Pomc* probe (#318511-C2) contained 17 oligonucleotide pairs and targeted region 21-921 (Acc. No. NM_139326.2) of the *Pomc* transcript. The *Th* probe (#314651-C2) contained 20 oligonucleotide pairs and targeted region 422-1403 (Acc. No. NM_012740.3) of the *Th* transcript. Negative and positive control probes were processed in parallel with the target probes to ensure RNA integrity and an optimal assay performance. On the day prior to the assay, the sections were mounted onto SuperFrost Plus slides (#631-9483; VWR, Radnor, PA, USA) and baked at 60 °C overnight in a HybEz oven (#321462). On the day of the assay, slides were first incubated for 7 min in hydrogen peroxide (#322335), submerged in Target Retrieval buffer (#322001) and rinsed in autoclaved Milli-Q purified water. The slides were quickly dehydrated in 100 % ethanol and allowed to air-dry. All the sections were then incubated with Protease Plus (#322331) for 30 min. The subsequent steps were performed according to the manufacturer’s protocol for the tyramide-based RNAscope® Multiplex Fluorescent v2 Assay (#323100). The *Agrp*, *Pomc* and *Th* probes were labelled with Opal 520 (1:500; FP1487A; PerkinElmer, Waltham, MA, United States), the *Ghsr* probe with Cy3 (1:2000; Akoya Biosciences, Menlo Park, CA, USA), and the *c-Fos* probe with Cy5 (1:3000; Akoya Biosciences). All the sections were counterstained with DAPI, coverslipped with ProLong® Diamond Antifade mountant (Thermo Fisher) and stored in the dark at 4 °C until imaging.

Images for the quantification of RNAscope data were captured using a laser scanning confocal microscope (LSM 700 inverted, Zeiss, Oberkochen, Germany) equipped with a Plan-Apochromat 40x/1.3 Oil DIC objective (used at the Centre for Cellular Imaging at Gothenburg University). Tile scans (5 × 5, 5 x 6 or 5 x 7 tiling, depending on the bregma) and Z-stacks (optical section of 1.0 μm) of the Arc-containing sections were captured unilaterally from rostral to caudal (2-4 sections per rat from 2.04 to 3.48 mm caudal to bregma). Laser intensities for the different channels were kept constant throughout the imaging process. The Z-stack images were processed using the maximum intensity projection function in the Zen Black software (Zeiss). The final images were then stitched and the cells counted in ImageJ/Fiji (NIH). The Cell counter plug-in was used to count positive cells and co-localization in the Arc. DAPI stain was used for cellular recognition. DAPI-identified cells with >1-3 dots/cell were defined as being positive for a given peptide. The quantification of the co-expression per hemisection was averaged for each brain and ultimately for each experimental group.

**Assessment of active ghrelin levels upon olfactory detection of PB**

Confirmation of the cue-induced feeding response

Another cohort of PB taste-familiar rats (n=21) was used to confirm the cue-induced hyperphagic effect. After habituation to PB taste, PB was encapsulated in the perforated balls and the intake of chow measured manually at 1, 3 and 6 hr post-cue introduction. The same time points were used for the control condition on the day before, in which spontaneous food intake was measured in the absence of PB odor.

Acyl-ghrelin assay

The same rats were re-exposed to either an olfactory PB cue-enriched environment (n=13) or to a non-enriched environment (perforated balls with empty tubes; n=8) for 1 hr, after which they were anesthetized and sacrificed to obtain blood for later assessment of active ghrelin. Following anesthesia with isoflurane and decapitation, trunk blood was immediately collected into EDTA-coated tubes containing 4-(2-Aminoethyl)benzenesulfonyl fluoride hydrochloride (AEBSF) to a final concentration of 1 mg/mL. We did not acidify the sample because it has been shown that HCl addition to AEBSF-treated samples does not provide enhanced hormone stability (7). To avoid cross-odor contamination, rats from different groups were kept in adjacent rooms. Sacrifices took place from 10 to 12 AM and groups were counterbalanced with respect to the time of sacrifice. Tubes were ultimately centrifuged to obtain plasma, which was aliquoted and stored at -80°C until processing. Plasma acyl-ghrelin levels were measured in duplicate using a commercial ELISA kit (#EZRGRA-90K; Merck Millipore, Darmstadt, Germany) following the manufacturer’s instructions. Samples were thawed only once.

**Statistics**

The program IBM SPSS Statistics 27 (IBM Corp., Armonk, NY, USA) was used for statistical analyses. Comparisons were carried out by one-way repeated measures ANOVA (rANOVA) when assessing the feeding response and meal patterns upon exposure to the olfactory PB cue (with conditioning [naïve, familiar] as “between factor” and cue [cue absent, olfactory PB cue, cue removal – when applicable] as “within factor” variables). We used paired samples *t*-tests to compare the cue-induced feeding response to that of the baseline measurement, as well as the intake of PB during the different taste conditionings in PB taste-familiar rats and mice, and a one-way ANOVA to assess the effects of the olfactory cue on active ghrelin levels. All other data were subjected to two-way ANOVA analyses (conditioning, cue). Given that we did not contemplate an explicit comparison between PB taste-naïve and familiar rats, additional paired samples *t*-tests or one-way ANOVAs (instead of any post hoc correction for multiple comparisons between the four groups) were used when significant main effects and/or interactions were obtained upon data split according to conditioning.

**Supplementary Results (statistical results and extended RNAscope results)**

**Olfactory detection of PB increases meal frequency to cause chow overconsumption in sated PB taste-familiar rats.**

One-way repeated measures analysis of variance (rANOVA) analyses revealed significant (or near-significant) effects of the PB olfactory cue on feeding measurements at 3 (F_[1,31]_=3.841, *p*=0.059), 6 (F_[1,31]_=6.922, *p*=0.013), 10 (F_[1,31]_=9.029, *p*=0.005) and 16 hr (F_[1,31]_=6.142, *p*=0.019), but not 1 or 24 hr (F_[1,31]_=1.345, *p*=0.255 and F_[1,31]_=0.003, *p*=0.958, respectively), as well as a conditioning x cue interaction for the 1 hr (F_[1,31]_=6.452, *p*=0.016) and 3 hr measurements (F_[1,31]_=5.898, *p*=0.021). After data split based on conditioning [naïve, familiar], paired samples *t*-tests for each of the time points studied demonstrated that PB taste-naïve animals did not change their food intake despite the presence of PB odor, neither acutely nor in the long-term, compared to their own baseline record (1 hr: t_[15]_=0.975, *p*=0.345; 3 hr: t_[15]_=0.344, *p*=0.735; 6 hr: t_[15]_=-1.400, *p*=0.182; 10 hr: t_[15]_=-1.598, *p*=0.131; 16 hr: t_[15]_=-0.824, *p*=0.423; 24 hr: t_[15]_=1.048, *p*=0.311) (Figure 1B). On the contrary, paired samples *t*-tests highlighted significant differences between the baseline and the cue exposure conditions in PB taste-familiar rats at 1 (t_[15]_=-2.618, *p*=0.019), 3 (t_[15]_=-2.994, *p*=0.009), 6 (t_[15]_=-2.242, *p*=0.041), 10 (t_[15]_=-2.684, *p*=0.017) and 16 hr (t_[15]_=-2.551, *p*=0.022), but not 24 hr (t_[15]_=-0.733, *p*=0.475) post-cue introduction in the home environment (Figure 1C). Cumulative 24-hr chow intake following cue removal was similar to that of the baseline and the cue settings in both PB taste-naïve and familiar rats (cue: F_[2,31]_=0.001, *p*=0.999, conditioning x cue: F_[2,31]_=0.712, *p*=0.499) (Figure S1).

Regarding meal patterns, rANOVA analyses for the entire 24-hr period revealed a significant effect of the olfactory cue on meal size (F_[1,31]_=4.164, *p*=0.050), but not meal frequency (F_[1,31]_=0.030, *p*=0.863), and a significant conditioning x cue interaction for meal frequency (F_[1,31]_=21.953, *p*<0.001). No significant differences were observed for the ingestion rate (cue: F_[1,31]_=0.168, *p*=0.684, conditioning x cue: F_[1,31]_=0.573, *p*=0.455): naïve rats, 2.35 ± 0.40 g/min vs. 1.93 ± 0.39 g/min (baseline vs. cue exposure, t_[15]_=0.686, *p*=0.503); familiar rats: 1.52 ± 0.24 g/min vs. 1.64 ± 0.40 g/min (baseline vs. cue exposure, t_[15]_=-0.330, *p*=0.746). After data split based on conditioning [naïve, familiar], paired samples *t*-tests showed that PB taste-naïve rats tended to increase the number of meals during the light phase upon exposure to the olfactory cue (t_[15]_=-2.055, *p*=0.058), although this effect was ultimately offset, since the same analysis confirmed reductions in the number of meal bouts during both the dark phase (t_[15]_=5.493, *p*<0.001) and the entire 24-hr period (t_[15]_=3.223, *p*=0.006) (Figure 1D). Paired samples *t*-tests also confirmed that this decrease in meal frequency was compensated by an increase in meal size during the dark phase (t_[15]_=-3.226, *p*=0.006) and the entire 24-hr period (t_[15]_=-2.133, *p*=0.050) (not significant for the light phase: t_[15]_=-0.804, *p*=0.434) (Figure 1F). For PB taste-familiar rats, paired samples *t*-tests highlighted significant (or near-significant) differences in meal frequency between the baseline and the cue exposure conditions during the light (t_[15]_=-2.298, *p*=0.036) and the dark phases (t_[15]_=-1.801, *p*=0.092), as well as for the entire 24-hr period (t_[15]_=-3.434, *p*=0.004) (Figure 1E). The analysis further confirmed that this effect was not compensated anyhow, because we could not detect any differences in meal size between the baseline and the cue exposure conditions during the light (t_[15]_=-0.835, *p*=0.417) or the dark phases (t_[15]_=0.640, *p*=0.532), or for the entire 24-hr period (t_[15]_=-0.427, *p*=0.676) (Figure 1G).

In a parallel study, a cohort of PB taste-familiar mice was used to determine whether olfactory detection of PB could likewise influence chow intake in mice. Paired samples *t*-tests between the control and the cue exposure conditions revealed that chow consumption in mice was not affected by the presence of PB odor in the home environment (1 hr: t[9]=-1.372, *p*=0.203; 3 hr: t[9]=-0.934, *p*=0.373; and 6 hr post-cue introduction: t[9]=-0.210, *p*=0.839) (Table S1).

Expectedly (if we consider a plausible neophobic response to a different taste and caloric content), PB consumption was greater during the second tasting session compared to the first one (rats: 87 % increase, t[15]=-6.299, *p*<0.001; mice: 67 % increase, t[9]=-3.191, *p*=0.011), and then plateaued during the last familiarization to PB taste (second vs. third PB tasting session in rats: t[15]=1.107, *p*=0.286; in mice: t[9]=-1.231, *p*=0.249) (Table S2). Hence, subjecting rodents to a single tasting experience to a palatable food does not seem optimal to foster its craving and subsequent seeking when cued by an olfactory stimulus.

**Olfactory detection of PB prompts food-seeking in sated PB taste-familiar rats.**

Two-way ANOVA analyses revealed significant (or near-significant) effects of the olfactory cue on the duration of the first contact with the perforated ball (F_[1,34]_=3.921, *p*=0.057) and the total time the rats spent in contact with the set-up (F_[1,34]_=15.660, *p*<0.001), but not on latency to approach the set-up (F_[1,34]_=1.953, *p*=0.172; Figure 2B). Further one-way ANOVAs (data split based on conditioning [naïve, familiar]) indicated that the duration of the first contact with the perforated ball set-up (F_[1,14]_=0.732, *p*=0.408) and the total time spent in contact with it (F_[1,14]_=2.459, *p*=0.141) were similar between the control and the olfactory PB-baited groups in PB taste-naïve rats (Figure 2C-D). On the contrary, both the duration of the first contact with the perforated ball set-up (F_[1,19]_=4.722, *p*=0.043) and the total time spent in contact with it (F_[1,19]_=18.229, *p*<0.001) did differ significantly between the control and the olfactory PB-baited groups within PB taste-familiar rats (Figure 2C-D).

**Olfactory detection of PB activates cells in the arcuate nucleus in sated PB taste-familiar rats.**

Two-way ANOVA analyses revealed a near-significant main effect of conditioning (F_[1,27]_=3.934, *p*=0.059), as well as a significant conditioning x cue interaction for the number of Fos-positive nuclei (F_[1,27]_=6.697, *p*=0.016). A one-way ANOVA analysis (data split based on conditioning [naïve, familiar]) indicated that the olfactory PB cue induced Fos activation in the Arc in PB taste-familiar rats (F_[1,12]_=4.988, *p*=0.047) (Figure 3D-E), but not in PB taste-naïve rats (F_[1,14]_=1.476, *p*=0.246) (Figure 3B-C).

**Intermingled and overlapping neuronal populations in the Arc are activated upon olfactory detection of PB in sated rats.**

Regarding triple co-expressions, a quarter of the total population of cells (25.3 ± 3.1%) co-expressing AgRP and GHSR were recruited upon olfactory detection of PB (i.e., were Fos+, Figure 4F). The majority of the AgRP cells activated by the PB cue (87.9 ± 1.8%) contained also GHSR, and almost half of the GHSR cells activated by the PB cue (46.0 ± 1.7%) were also positive for AgRP (Figure 4F). More than a third of the total population of cells (39.4 ± 3.8%) co-expressing POMC and GHSR were recruited upon olfactory detection of PB (Figure 5F). Similarly, more than a third of the POMC cells activated by the PB cue (38.0 ± 8.8%) contained also GHSR; and nearly a third of the GHSR cells that were activated by the PB cue (28.6 ± 1.8%) were also positive for POMC (Figure 5F). Finally, approximately a quarter of the total population of cells (27.8 ± 7.3%) co-expressing TH and GHSR were recruited upon olfactory detection of PB (Figure 6F). Three quarters of the TH cells activated by the PB cue (77.9 ± 5.3%) contained also GHSR; and almost third of the GHSR cells activated by the PB cue (30.4 ± 2.2%) were also positive for TH (Figure 6F).

On the other hand, over half of GHSR+ cells (56.5 ± 1.9%) co-expressed AgRP and 83.2 ± 4.0% of AgRP cells expressed GHSR (Figure 4D); 14.5 ± 1.8% of GHSR+ cells co-expressed POMC and, interestingly, approximately a third of POMC neurons (32.2 ± 4.2%) were GHSR+ (Figure 5D); and finally, 20.1 ± 2.0% of GHSR+ cells expressed TH and two thirds of TH cells (68.0 ± 2.0%) expressed GHSR (Figure 6D).

**Olfactory detection of PB triggers release of active ghrelin in sated PB taste-familiar rats.**

To confirm the food intake data, we carried out manual food intake measurements upon exposure to the olfactory PB cue in PB taste-familiar rats. Paired samples *t*-tests pointed to significant differences in food intake between the control and the cue exposure conditions 1 (t_[20]_=-2.951, *p*=0.008) and 6 hr (t_[20]_=-2.769, *p*=0.012), but not 3 hr (t_[20]_=-0.997, *p*=0.331), post-cue introduction in the home environment (Figure 7A). Circulating levels of active ghrelin were likewise increased in sated PB taste-familiar rats upon 1-hr exposure to an olfactory PB cue, as indicated by one-way ANOVA (F_[1,20]_=4.497, *p*=0.047) (Figure 7B).

**Supplementary Tables**

| **Table S1.** Cumulative chow consumption (1, 3 and 6 hr post-cue presentation) in mice presented an olfactory PB cue in the home environment | | | |
| --- | --- | --- | --- |
|  | **1-hr chow intake (g)** | **3-hr chow intake (g)** | **6-hr chow intake (g)** |
| **Non-enriched environment**  (perforated balls with empty tubes) | 0.044 ± 0.025 | 0.167 ± 0.04 | 0.363 ± 0.073 |
| **Olfactory PB cue-enriched environment**  (perforated balls with PB-filled tubes) | 0.137 ± 0.056 | 0.23 ± 0.051 | 0.388 ± 0.083 |
| PB, peanut butter; n=10. No significant differences were found. | | | |

| **Table S2.** Intake of PB during the 1-hr lasting PB tasting conditionings in both rats and mice | | | |
| --- | --- | --- | --- |
|  | **1-hr PB intake (g)**  **Day 1** | **1-hr PB intake (g)**  **Day 2** | **1-hr PB intake (g)**  **Day 3** |
| **Rats** | 2.01 ± 0.246 | 3.758 ± 0.268 *** | 3.49 ± 0.3 |
| **Mice** | 0.297 ± 0.056 | 0.497 ± 0.074 * | 0.56 ± 0.088 |
| PB, peanut butter; rats, n=16; mice, n=10. Symbols indicate significant differences vs. the first PB tasting session at * *p* < 0.05 or *** *p* < 0.001 | | | |

| **Table S3.** Acoustic measurements of the interaction with the perforated ball set-up in PB taste-naïve and PB taste-familiar rats | | | | |
| --- | --- | --- | --- | --- |
|  | | **Minimum (dB)** | **Average (dB)** | **Maximum (dB)** |
| **PB taste-naïve rats**  **(n=15)** | No cue | 26 | 32 | 55 |
|  | PB cue | 30 | 42 | 61 |
|  | ∆ | **4** | **10** | **6** |
| **PB taste-familiar rats**  **(n=20)** | No cue | 29 | 37 | 58 |
|  | PB cue | 34 | 50 | 68 |
|  | ∆ | **5** | **13** | **10** |
| **Percentage difference (%∆)** | | **25** | **30** | **67** |
| PB, peanut butter | | | | |

**Supplementary Figure**

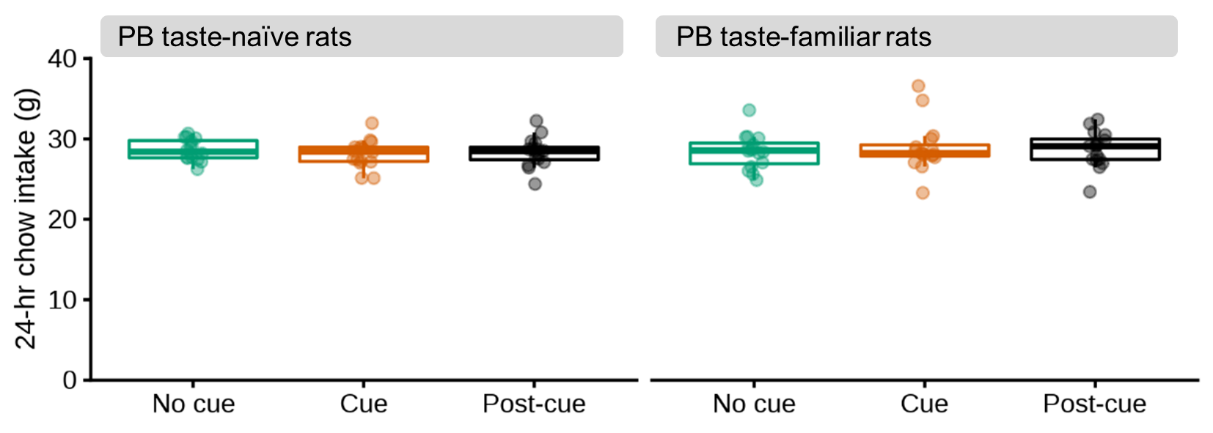

**Figure S1. Comparison of the cumulative intake of chow within 24 hr following cue removal with that of the baseline and PB cue settings.**

24-hr cumulative chow intake (g) during the baseline (green), PB cue (orange) and post-cue (black) settings in PB taste-naïve rats (left, n=16) and PB taste-familiar rats (right, n=16) Cumulative 24-hr chow intake following cue removal was similar to that of the baseline and the cue settings in both PB taste-naïve and familiar rats. In all cases, the thick line corresponds to the median, boxes show first and third quartiles and whiskers represent minimum and maximum values.

**Supplementary References**

1.	Stoltenborg I, Peris-Sampedro F, Le May MV, Bake T, Schele E, Adan RA, et al. (2020): Primary food cues engage pathways involved in over-eating and reward-seeking in rats. *European Neuropsychopharmacology*, 28 January, 2020 ed, pp S 36-S 37.

2.	Boggiano MM, Dorsey JR, Thomas JM, Murdaugh DL (2009): The Pavlovian power of palatable food: lessons for weight-loss adherence from a new rodent model of cue-induced overeating. *International journal of obesity (2005)*. 33:693-701.

3.	Farley C, Cook JA, Spar BD, Austin TM, Kowalski TJ (2003): Meal pattern analysis of diet-induced obesity in susceptible and resistant rats. *Obesity research*. 11:845-851.

4.	Crossley E, Biggs T, Brown P, Singh T (2021): The Accuracy of iPhone Applications to Monitor Environmental Noise Levels. *The Laryngoscope*. 131:E59-e62.

5.	Lockie SH, McAuley CV, Rawlinson S, Guiney N, Andrews ZB (2017): Food Seeking in a Risky Environment: A Method for Evaluating Risk and Reward Value in Food Seeking and Consumption in Mice. *Frontiers in neuroscience*. 11:24.

6.	Peris-Sampedro F, Stoltenborg I, Le May MV, Zigman JM, Adan RAH, Dickson SL (2021): Genetic deletion of the ghrelin receptor (GHSR) impairs growth and blunts endocrine response to fasting in Ghsr-IRES-Cre mice. *Molecular metabolism*. 51:101223.

7.	Blatnik M, Soderstrom CI (2011): A practical guide for the stabilization of acylghrelin in human blood collections. *Clinical endocrinology*. 74:325-331.
